# SWI/SNF remains localized to chromatin in the presence of *SCHLAP1*

**DOI:** 10.1101/322065

**Authors:** Jesse R. Raab, Keriayn N. Smith, Camarie C. Spear, Carl J. Manner, J. Mauro Calabrese, Terry Magnuson

## Abstract

SCHLAP1 is a long-noncoding RNA that is prognostic for progression to metastatic prostate cancer and promotes an invasive phenotype. SCHLAP1 is reported to function by depleting the core SWI/SNF subunit, SMARCB1, from the genome. SWI/SNF is a large, multi-subunit, chromatin remodeling complex that can be combinatorially assembled to yield hundreds to thousands of distinct complexes. Here, we investigated the hypothesis that SCHLAP1 affects only specific forms of SWI/SNF and that the remaining SWI/SNF complexes were important for the increased invasion in SCHLAP1 expressing prostate cells. Using several assays we found that SWI/SNF is not depleted from the genome by SCHLAP1 expression. We find that SCHLAP1 induces changes to chromatin openness but is not sufficient to drive changes in histone modifications. Additionally, we show that SWI/SNF binds many coding and non-coding RNAs. Together these results suggest that SCHLAP1 has roles independent of canonical SWI/SNF and that SWI/SNF broadly interacts with RNA.

## Main

The long non coding RNA (lncRNA) Second Chromosome Locus Associated with Prostate cancer 1 (*SCHLAP1*) is prognostic for metastatic disease and is a promising biomarker in prostate cancer ^1,2^. *SCHLAP1* is proposed to function by antagonizing the SWI/SNF complex through direct interaction leading to complete disruption of SWI/SNF genomic occupancy ^1^. Evidence for this mechanism comes from the reported loss of SMARCB1 occupancy measured by ChIP-seq ^1^. SWI/SNF is a large multi-subunit chromatin remodeling complex that can be combinatorially assembled to yield hundreds to thousands of biochemically distinct complexes, each of which may have unique functions ^3–6^. We investigated the alternate hypothesis that distinct forms of SWI/SNF are affected in disparate ways through *SCHLAP1* expression. However, a variety of biochemical and genomics assays we demonstrate that SWI/SNF occupancy is unaffected by *SCHLAP1* expression, in contrast to previously reported results. *SCHLAP1* expression induces changes to chromatin state measured by ATAC-seq, but does not lead to changes in histone modification states. Using RNA immunoprecipitation we show that SWI/SNF binds both coding and non-coding RNA, raising the possibility that the reported *SCHLAP1* SWI/SNF interaction is due a broad interaction between RNA and SWI/SNF and that *SCHLAP1* may function through another mechanism.

We hypothesized that SCHLAP1 expression affected the composition and occupancy of SWI/SNF complexes in prostate cancer. We first validated the reported interaction between SMARCB1 and *SCHLAP1* using 22Rv1 and LNCaP prostate cancer cells. Consistent with the previous report we observed an interaction between SMARCB1 and SCHLAP1 (Fig. 1A). We next wished to study SWI/SNF dynamics in a controlled model system. To do this we generated SCHLAP1 over-expressing benign prostate epithelial cells (RWPE1;*SCHLAP1*) or control cells RWPE1;*LACZ*)^1^ (*SCHLAP1* construct gift of A. Chinnayian). This model is the same one used to originally suggest global depletion of SMARCB1 by SCHLAP1 ^1^. Similar to the results of Prensner et. al., we saw strong induction of *SCHLAP1* in these cells upon overexpression, and we did not observe any changes in SWI/SNF subunit expression at the protein level (Fig 1B, Supp. Fig. 1A). Expression of *SCHLAP1* did not induce changes to proliferation in two-dimensional culture, and did not affect the ability of RWPE1 cells to grow from low cell numbers in low-adherent plates (Supp. Fig. 1B, C), consistent with the previous reports. Finally, we assessed the functional change demonstrated previously by performing invasion assays on RWPE1;*LACZ* and RWPE1;*SCHLAP1.* This confirmed an increase in invasion by cells expressing *SCHLAP1* (Fig 1C).

**Figure 1.**
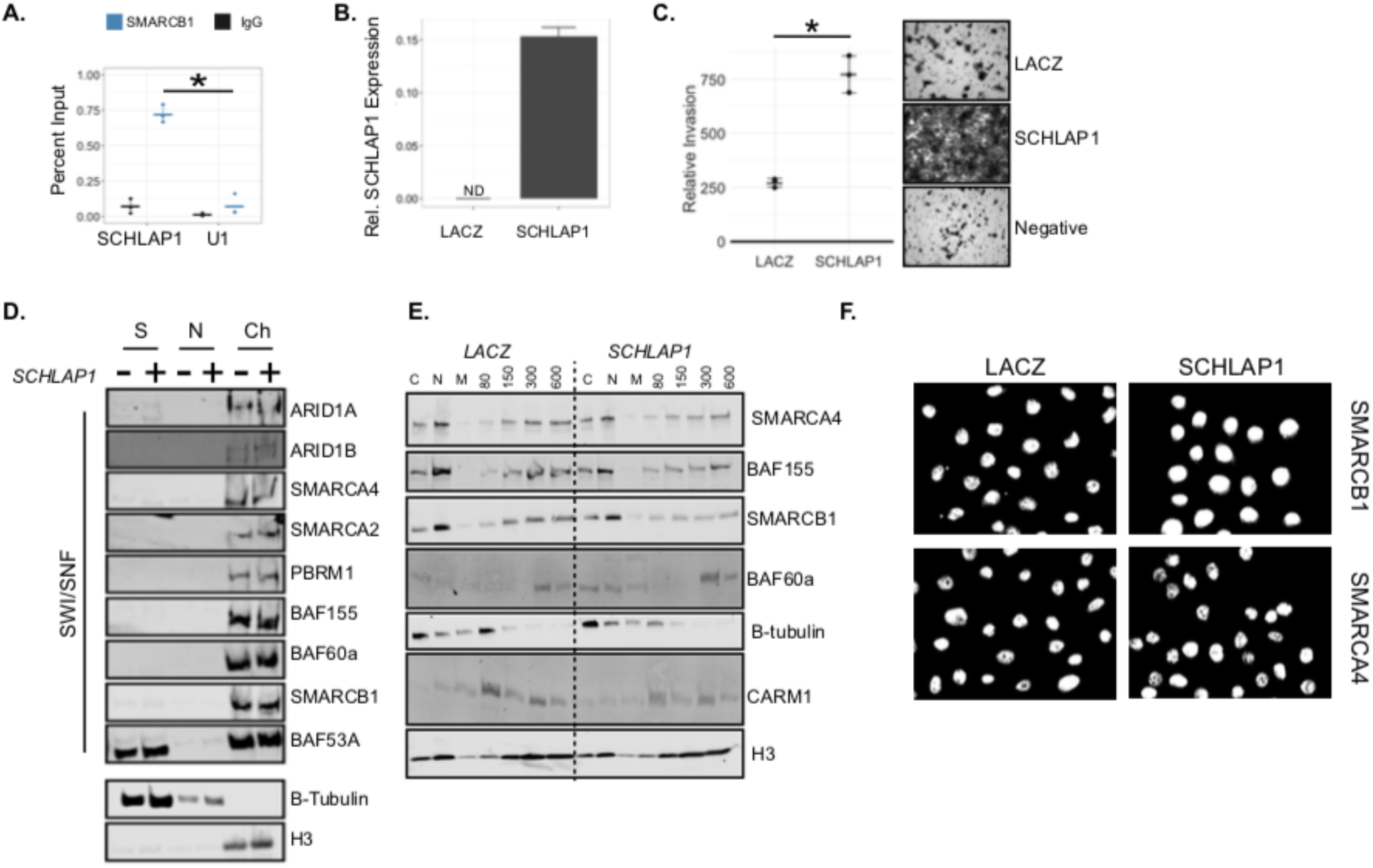
SCHLAP1 does not evict SWI/SNF from chromatin. A. Native RIP for SMARCB1, asterisk denotes p-value < 0.05 (t-test) between *SCHLAP1* and U1 primer sets for SMARCB1 IP, n = 3). B. SCHLAP1 expression in RWPE1;*LACZ* and RWPE1;*SCHLAP1* cells quantitative relative to GAPDH, n = 3. C. Invasion assay of RWPE1;*LACZ* and RWPE1;*SCHLAP1* cells quantitative by fluorescent intensity (Licor). Representative from n = 3 invasion assays, p-value < 0.05 (t-test). D. Chromatin fractionation showing SWI/SNF subunit presence in soluble (S), nuclear (N), or Chromatin (Ch) fractions from RWPE1;*LACZ* and RWPE1;*SCHLAP1* cells. E. Salt fractionation of RWPE1;*LACZ* and RWPE1;*SCHLAP1* cells for SWI/SNF subunits from whole cell (C), nuclei (N), micrococcal nuclease digestion (M) or salt fraction (mM). F. Immunofluorescence of RWPE1x;*LACZ* or RWPE1;*SCHLAP1* cells SMARCB1 or SMARCA4.

Together these results show our model faithfully recapitulates the previous functional aspects of *SCHLAP1* overexpression in RWPE1 cells and we validated the physical interaction between *SCHLAP1* and SMARCB1^1^.

To investigate which SWI/SNF subunits were depleted from chromatin upon *SCHLAP1* expression, we fractionated the RWPE1;*LACZ* and RWPE1; *SCHLAP1* cells into cytosolic, soluble nuclear, and insoluble chromatin enriched fractions ^7^. Surprisingly, all SWI/SNF subunits assayed remained strongly enriched in the chromatin fraction and we observed no gross differences between RWPE1;*LACZ* and RWPE1; *SCHLAP1* (Fig. 1D). We tested the association with chromatin using an orthologous biochemical method by using varying concentrations of salt to extract chromatin associated proteins ^8^. Consistent with our fractionation experiments, SWI/SNF subunits in both cell lines were extracted from nuclei under higher salt conditions in fractions containing histone H3 in both RWPE1;*Lacz* and RWPE1;*SCHLAP1* cells. SWI/SNF localized to distinct salt fractions compared to the non-chromatin associated protein β-tubulin and the arginine methyltransferase CARM1 (Fig. 1E). Consistent with these biochemical experiments, we found that SMARCA4 and SMARCB1 localization was not affected in RWPE1;*SCHLAP1* cells by immunofluorescence (Fig. 1F). Next, we performed co-immunoprecipitation for SMARCA4 and SMARCB1 and found that several SWI/SNF complex members could be immunoprecipitated together (Supp. Fig 1D). This demonstrates that the SWI/SNF complex remains intact in the presence of *SCHLAP1.* Finally, we used a malignant rhabdoid tumor cell line where SMARCB1 can be induced by culturing cells with doxycycline (Kind gift of B. Weissman).

The expression of SMARCB1 in these cells causes growth arrest ^9^. We reasoned that if *SCHLAP1* disrupted SMARCB1 chromatin occupancy, over-expression of *SCHLAP1* should allow G401 to continue growing following induction of SMARCB1. However, SMARCB1 induction led to growth arrest in a dose dependent manner in both cell lines (Supp. Fig. 2A-C). Together these results demonstrate that *SCHLAP1* does not induce changes to SWI/SNF composition or association with chromatin.

To validate these biochemical results we performed ChIP-seq for three SWI/SNF subunits in RWPE1; *SCHLAP1* cells. We used the core subunit SMARCB1, which was previously assayed by Prensner et al., as well as both ATPase subunits, SMARCA2 and SMARCA4. In contrast to the previous report, we identify robust binding for all three subunits in RWPE1 cells expressing *SCHLAP1* (Fig. 2A). In RWPE1; *SCHLAP1* cells we identified 6490, 22185, and 51505 peaks for SMARCB1, SMARCA2, and SMARCA4 respectively (Supplemental Table 1). This large number of peaks is in contrast to the original report which identified ~6500 SMARCB1 peaks in RWPE1; *LACZ* cells and nearly zero peaks in the *SCHLAP1* expressing cells. The numbers of peaks are consistent with previous work from our lab showing 30,000-45,000 SMARCA4 peaks ^10^. Additionally we and others have reported a large number of SWI/SNF peaks for a variety of subunits ^3,10–12^. The vast majority of SMARCA2 peaks overlapped a SMARCA4 peak, consistent with our previous results ^10^ (Fig. 2B). The differences in numbers of peaks between subunits is likely due to differences in antibody efficiency, or subunits that are less accessible to DNA and cross linking relative to the rest of the complex.

**Figure 2.**
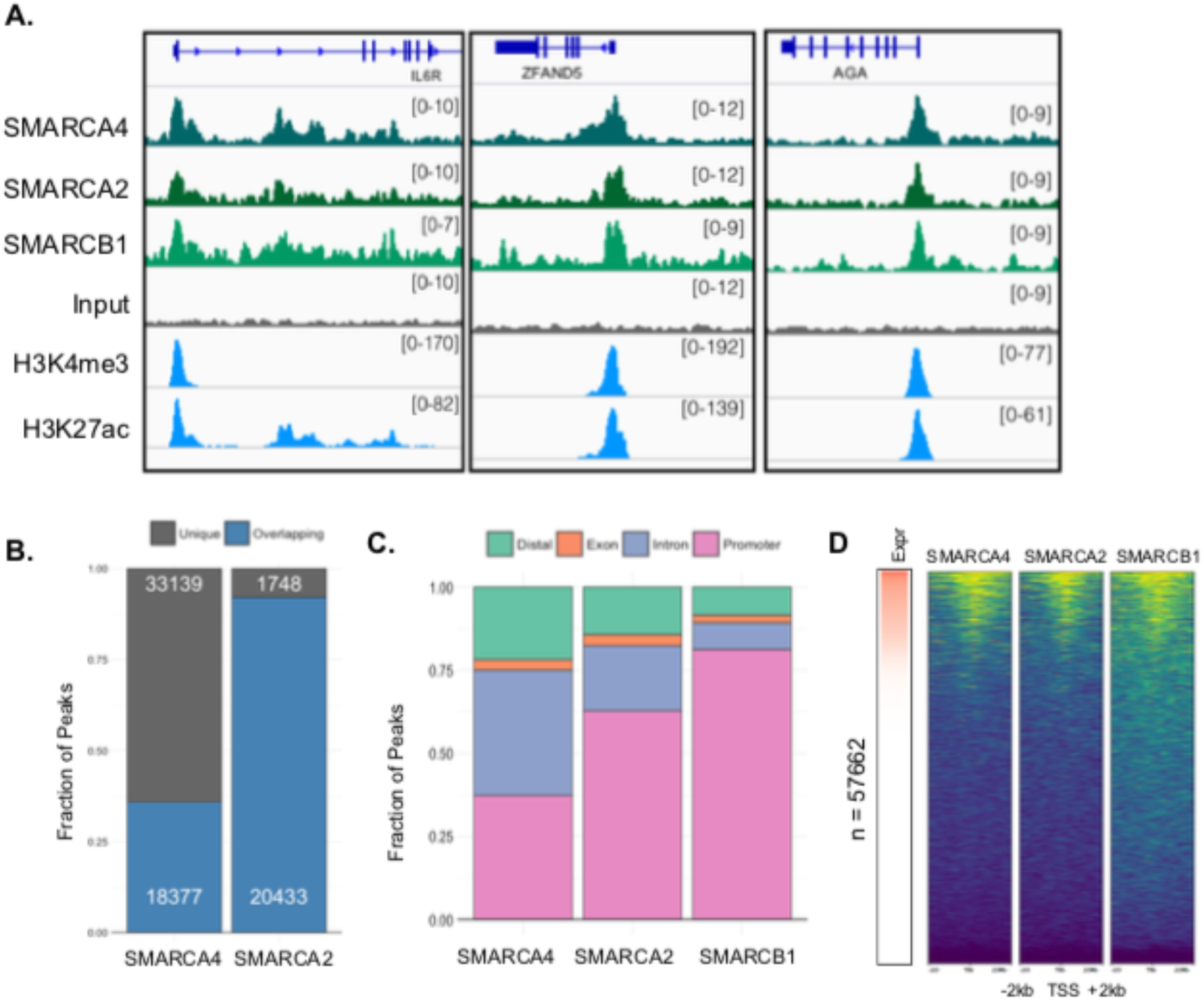
SWI/SNF occupancy in the presence of *SCHLAP1.* A. Example browser images for SMARCA4, SMARCA2, and SMARCB1 from RWPE1;*SCHLAP1* cells ChIP-seq experiments. B. Overlap between SMARCA2 and SMARCA4 peaks. C. Genomic location of peaks. D. Occupancy for SMARCA4, SMARCA2, and SMARCB1, centered on all transcription start sites (GENCODE), aligned by expression in RWPE1 cells.

**Fig. 3.**
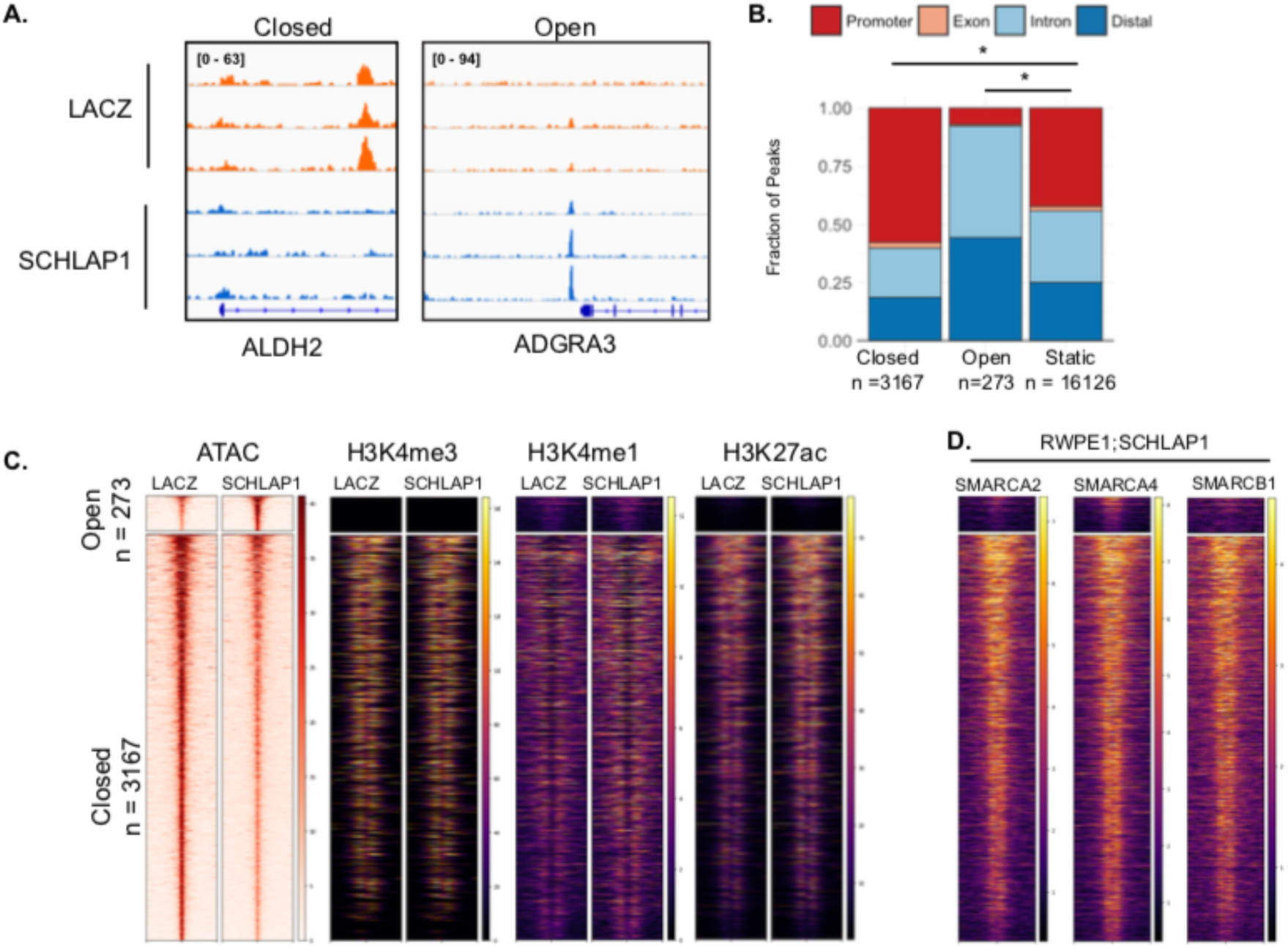
*SCHLAP1* induces open chromatin changes but does not alter histone modifications. A. Example loci representing open and closed chromatin regions from three replicate ATAC-seq experiments. B. Genomic loci associated with open, closed, and static sites. Asterisk denotes p-value < 0.01, chi squared test. C. Heatmap displays open chromatin, H3K4me3, H3K4me1, or H3K27ac signal in RWPE1;*LACZ* or RWPE;*SCHLAP1.* Rows are ordered according to level of open chromatin signal. D. Heatmap shows the level of three SWI/SNF subunits present in *RWPE1*;*SCHLAP1* at altered open chromatin sites. Order is the same as in panel C.

SWI/SNF peaks were predominantly located at promoters (45%-75% Fig. 2C), with SMARCA4 having the most non-promoter peaks (~55% non-promoter). We monitored signal across all annotated transcription start sites in GENCODE and identified concordant strong signal across approximately 25% of TSS. SWI/SNF binding was most prominent at highly expressed genes with little to no occupancy at non-expressed genes (Fig. 2D, GSE98898 for expression data)^13^.

These results demonstrate that *SCHLAP1* does not function by disrupting SWI/SNF occupancy genome-wide and raises the question of how *SCHLAP1* functions to promote invasion and progression to metastatic disease. To determine if *SCHLAP1* expression induced chromatin changes, we performed ATAC-seq on RWPE1;*LACZ* and RWPE1;*SCHLAP1* cells ^14^ We identified 273 and 3167 sites that open and close respectively (Supplemental Table 2). The sites that opened were more likely to be located distally or in introns of genes (Figure 3A, B). Sites that open upon *SCHLAP1* expression were enriched for distinct motifs from those that close (Supp. Fig 3A, Supplemental Table 3, 4). Open sites were enriched in motifs for TEAD and AP1 transcription factors. AP1 and TEAD motifs were closely spaced consistent with a potential role in oncogenic enhancers ^15–17^. To test if these sites became activated enhancers, we performed ChIP-seq for H3K4me1, H3K4me3, and H3K27ac in RWPE1;*LACZ* and RWPE1; *SCHLAP1* cells (Fig. 3C). No major differences were identified when comparing ChIP-seq signaling between RWPE1;*LACZ* and RWPE1; *SCHLAP1* cells. We looked at whether SWI/SNF subunits were localized to the dynamic ATAC sites. SMARCA2, SMARCA4, and SMARCB1 were more highly enriched in RWPE1; *SCHLAP1* on sites that lose chromatin accessibility suggesting that SWI/SNF remains occupied at these sites in the presence of *SCHLAP1.* GO analysis of the genes associated with open sites revealed pathways involved in responses to NFKB signaling, epithelial to mesenchymal transitions, and nucleotide metabolism (Supp. Fig 3B). Genes associated with closed sites were enriched for pathways involved in cell adhesion and signal transduction via RAS, MAPK, and WNT (Supp. Fig 3B). We used microarray data from Prensner et al. to investigate if closed or open chromatin sites were associated with gene expression changes, but did not detect a significant association between gene expression changes and the direction of altered chromatin openness (Supp. Fig. 4A. 4B)^1^.

As *SCHLAP1* did not exert broad effects on SWI/SNF occupancy, we next investigated whether SWI/SNF interacts with other lncRNA. Numerous reports suggest a role for lncRNA-SWI/SNF interactions ^18–23^. We performed crosslinked RIP-seq for SMARCA4 and a general splicing factor that would not be expected to have a similar interactions as a chromatin remodeler (Splicing Factor Proline And Glutamine Rich - *SFPQ*) in 22Rv1 cells in LNCaP cells (Fig. 4A) ^24^. Consistent with previous observations of chromatin regulators, we saw widespread enrichment of SMARCA4 on most expressed transcripts relative to an IgG control (Supplemental Table 5, 6) ^24^ The pattern of SWI/SNF enrichment appeared relatively uniform throughout the transcript, including introns, suggesting SMARCA4 associates frequently with primary transcripts (Fig 4C-F). In contrast, SFPQ had more restricted binding patterns, with most enrichment on exons and high levels of enrichment at the 3’ UTR of many transcripts. We observed SMARCA4 and SFPQ signal at *SCHLAP1,* although the level of enrichment was not strong or markedly different between the proteins. Among the enriched transcripts was *NEAT1,* which is a known interacting partner of both SWI/SNF ^19^ and SFPQ ^25,26^. SFPQ is believed to play a role in alternate isoform usage of *NEAT1* through regulating 3’ end processing, and its binding pattern at SFPQ reflects a role on both the short and long isoforms (Fig. 4A, short arrow)^27^. Additionally, we identified high levels of SMARCA4 associated with MALAT1, while SFPQ was lower at this gene (Fig 4A,B). A recent report showed a functional interaction between SMARCA4, HDAC9, and *MALAT1,* further supporting the specificity of our result ^23^. The broad pattern of enrichment suggests SWI/SNF may have a frequent interaction with RNA that might not be sequence specific.

**Fig 4.**
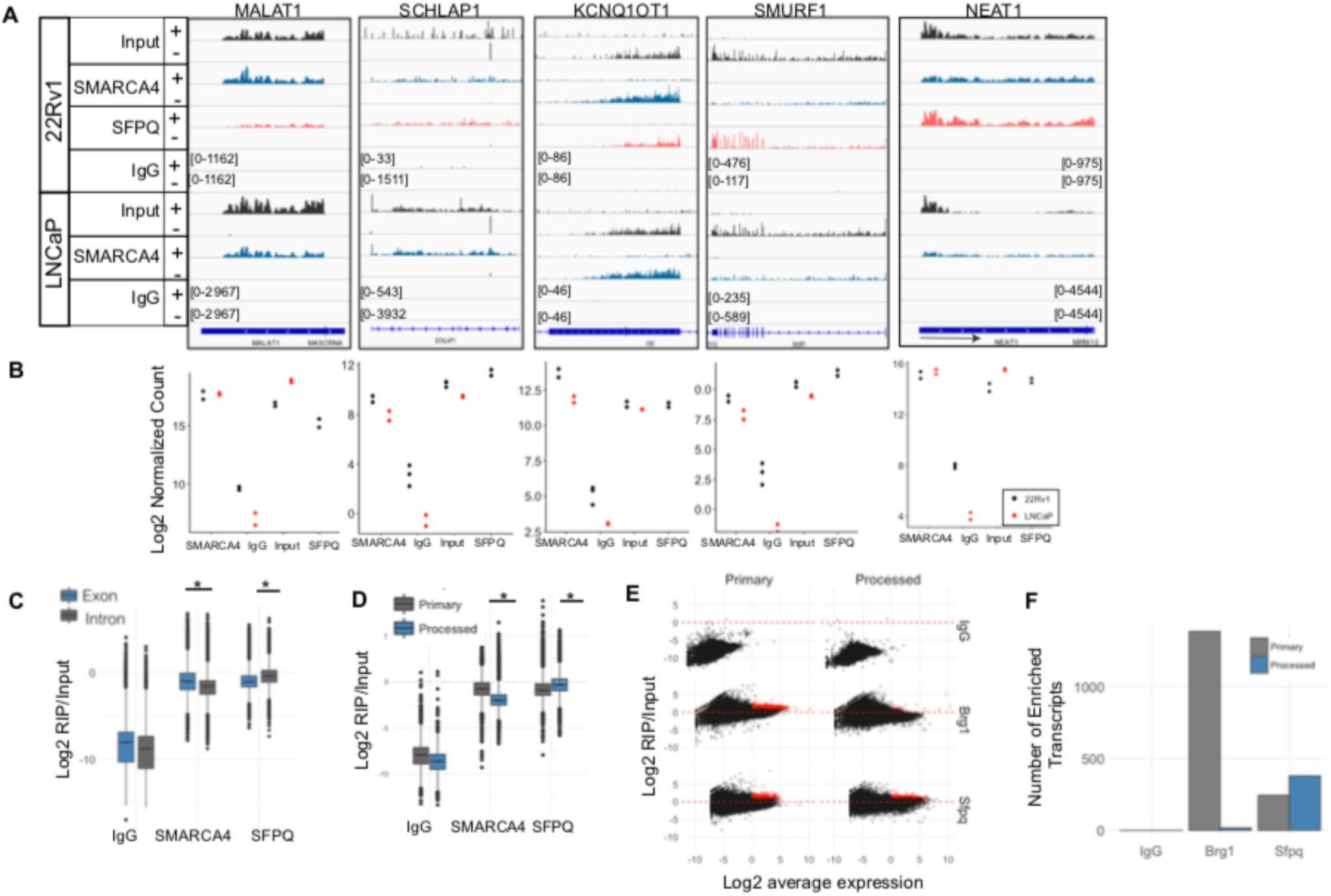
SMARCA4 binds many RNAs. A. Example loci associated with SMARCA4 or SFPQ in 22Rv1 and LNCaP cells. Strand specific RNA is shown (+/-) at each loci. Within cell lines and on the same strand scales are equal. Short isoform of *NEAT1* is denoted by an arrow in the gene track. B. Quantitation of signal at each of the genes in panels A for each antibody and each cell line. C. Enrichment of reads mapping to exons compared to introns for each IP, asterisk denotes p-value < 2.2e-16, wilcoxon test. D. Enrichment of reads mapping to primary compared to processed transcripts, asterisk denotes p-value < 2.2e-16, wilcoxon test. E. Log2 fold change relative to input is plotted against the average expression of a transcript for both the primary and processed transcripts for the three antibodies tested. Red points indicate those genes greater than log2 fold change of 1 and an average log2 fold change greater than 0. F. Number of transcripts assigned to the red points in panel E.

Together these results demonstrate that the interaction between SWI/SNF and *SCHLAP1* does not lead to a global depletion of SWI/SNF from the genome as previously reported ^1^. However, we confirm that *SCHLAP1* induces RWPE1 cells to become more invasive, and our results suggest that this phenotype may be driven by a SWI/SNF independent mechanism. As *SCHLAP1* has shown prognostic value for identifying metastatic prostate cancer, it is important to identify the mechanism by which it promotes invasion ^2^. *SCHLAP1* expression induces moderate changes to chromatin but appear insufficient to drive changes to histone modifications (Fig. 3). Our RNA immunoprecipitation results suggest that SWI/SNF interactions with RNA may be widespread, as reported for other chromatin regulators ^24^. *In vitro,* RNA is capable of inhibiting chromatin remodeling activity and this may provide a mechanism of regulating chromatin remodeling activity via locus-specific broad SWI/SNF-RNA interactions ^22^. Future studies are needed to improve our understanding of how widespread interactions between chromatin remodelers and RNA affect chromatin and transcription.

## Methods

### Cell Culture

RWPE1 cells were purchased from ATCC (CRL-11609, Lot 61840713) and grown in Keratinocyte Serum Free Medium (Life Technologies) supplemented with penicillin and streptomycin. Cells were STR profiled by ATCC before purchase, and all experiments were performed on low passage cells (< 10 passages). Lenti6-*SCHLAP1* was a kind gift of A. Chinnayian ^1^. Lenti6-Lacz was generated by removing SCHLAP1 from the above plasmid and cloning LACZ from pLentipuro-LACZ into pLenti6 from which SChLAP1 was removed by EcoRI-BamHI digest. Lentiviral particles were produced using psPAX2 and pMD2.G to package plasmids in 293T cells. RWPE1;*Lacz* and *RWPE1*;*SCHLAP1* cells were then generated by transduction with lentivirus and selecting with 2.5ug/mL blasticidin for 1 week. 22RV1 and LNCaP Cells were purchased from ATCC (CRL-2505, Lot 60437301; CRL-1740, Lot 62129998) and grown in RPMI 1640 with 10% Fetal Bovine Serum supplemented with penicillin and streptomycin. Cells were STR profiled by ATCC prior to purchase and were used at early passages (< 10 passages). All cells were tested and free of mycoplasma contamination.

### Antibodies

SMARCA2 (Cell Signaling 11966), SMARCA4 (Abcam ab110641), SMARCB1 (Abcam ab192864), BAF53A (Abcam ab3882), BAF60A (BD Biosciences 611728), BAF155 (Cell Signaling D7F8S), BAF180 (Bethyl A301-591A), SFPQ (Genetex GTX114209), NCL (Bethyl A300-711A), ARID1A (Abcam ab182560), ARID1B (Abcam ab57461), ARID2 (ThermoFisher PA5-35857), H3K4me3 (Abcam ab8580), H3K4me1 (Active Motif 39635), H3K27ac (Active Motif 39133), H3 (epicypher 13-0001)

### Invasion/Migration Assay

Invasion was assessed using a Corning BioCoat Matrigel Invasion Assay with 8μm pore size. Cells were seeded into Boyden Chambers at 100,000 cells/well in 100μl of normal growth media. The lower chamber was filled with either normal growth media supplemented with 10% FBS or PBS as a negative control. Cells were allowed to invade for 40.25 hours after which the top chamber was swabbed to remove non-invading cells. Invading cells were fixed and stained with crystal violet. Membranes were imaged using a Zeiss Axio Imager for representative images or scanned using a Licor fluorescent imager for quantitation. Results are reported as the average of three biological replicates. For each replicate, crystal violet fluorescence was measured and background fluorescence was subtracted. Background fluorescence of the membranes was obtained from a negative control.

### RNA Immunoprecipitation

RIP experiments were performed using a modified version of a protocol from Hendrickson et al.. ^24^ ***Fixation*** *-* 22Rv1 and LNCaP cells were fixed in 0.3% methanol free formaldehyde for 30 minutes at 4°C. Formaldehyde was quenched with 125mM glycine for 5 minutes at room temperature. Plates were washed 3 times in room temperature PBS and cells were scraped in 1mM PMSF in PBS. Cells were snap frozen in liquid nitrogen and stored at -80°C.

***Fragmentation and IP*** *-* Cells were resuspended in 0.5mL RIPA buffer (50mM Tris-HCl pH 8, 1% Triton X-100, 0.5% sodium deoxycholate, 0.1% SDS, 5mM EDTA, 150mM KCl) + 0.5mM DTT with 1X protease inhibitors (Sigma) and 2.5uL RNAsin and incubated on ice for 10 minutes prior to lysing using a Bioruptor (Diagenode) for two cycles of 30 seconds on and 1 minute off, followed by centrifugation at 4°C for 10 minutes at max speed. Protein A magnetic beads (Biorad,) were pre-conjugated with antibody for 2 hours at 4 degrees. Cells were then incubated overnight at 4 degrees with antibody conjugated beads. Beads were washed consecutively with fRIP buffer (25mM Trix-HCl pH 7.5, 5mM EDTA, 0.5% Ipegal CA-630, 150mM KCl), 3 times in ChIP buffer (50mM Tris-HCl pH 7.5, 140mM NaCl, 1mM EDTA, 1mM EGTA, 1% Triton X-100, 0.1% sodium deoxycholate, 0.1% SDS), 1x in high salt buffer (ChIP buffer, but with 500mM NaCl) and 1x in fRIPbuffer. All washes were performed for 5 minutes at 4 degrees. After final wash, beads were resuspended in 56uL water and 33uL of 3X reverse crosslinking buffer (3x PBS, 6% N-lauroyl sarcosine, 30MM EDTA) supplemented with 5mM DTT, add 20uL proteinase K, and 1uL RNAsin and incubated 1 hour at 42 degrees, 1 hour at 55 degrees, and 30 minutes at 65 degrees. Following elution and crosslink reversal, trizol was used to extract RNA. Finally, RNA in the aqueous phase was supplemented with 1 volume of ethanol and purified using a Zymo-spin IC column, including the on-column DNAse digestion, per the manufacturer's instruction. RNA was eluted in 15uL ddH20 and used to prepare RNA-seq libraries or to synthesize cDNA for qpCR.

RNA-seq libraries were prepared by using equal volumes of IPs and included 1uL of 1:250uL dilution of ERCC spike-in mix 1 (Life Technologies). Input libraries were prepared from the same amount of RNA as the IP with the most RNA, and spike-ins were added as above. Libraries were then prepared using the Kapa Ribo-zero kit per manufacturers instructions, pooled, and sequenced on using single-end 75bp reads on a Nextseq 500.

### RIP-seq Analysis

Reads were aligned using STAR to hg38 with gencode v27 annotations that included ERCC spike-ins and using --quantMode GeneCounts. In parallel an alignment-free quantitation method (Salmon version 0.8) was used to quantitate against gencode v27 containing ERCC spike ins and an extra transcript for each gene that contained the whole genomic DNA locus to represent an unspliced transcript. Browser tracks were generated by converting BAM files to bigWig files using Deeptools (v 2.5.2) and scaling tracks according to the 75% percentile of the ERCC-spike ins. Enrichment relative to IgG was calculated using DESeq2 with sizeFactors generated from only the ERCC spike-in data. For visualization relative to Input in Figure 4C,D,E normalized count values were used and log2 (RIP/Input) was calculated for each gene. RIP-seq data are available under GEO accession GSE114393

### Chromatin Fractionation

Chromatin fractionation was performed as previously described ^7^. Approximately 1e7 cells were washed and resuspended in 200uL Buffer A (10mM Hepes pH 7.9, 10mM KCl, 1.5mM MgCl2, 0.34M Sucrose, 10% Glycerol, 1mM DTT, supplemented with 1X protease inhibitors and 1mM PMSF). Triton X-100 was then added to 0.1% final concentration and cells were incubated for 8 minutes on ice. Cells were then centrifuged at 1300g x 5 minutes at 4 degrees. The supernatant was saved as the cytosolic fraction and the pellet (nuclei) was washed once with Buffer A. Cells were lysed for 30 minutes on ice in 100uL Buffer B (3mM EDTA, 0.2MM EGTA, 1mM DTT) supplemented with 1X protease inhibitors and 1mM PMSF and centrifuge at 1700g x 5 minutes at 4 degrees. The supernatant was saved as the nucleoplasmic fraction and the insoluble material was resuspended in 1X laemmli buffer, and sonicated twice for 30 seconds on high power (Bioruptor). All lysates were boiled for 10 minutes in 1X laemmli buffer with 0.1M DTT and used in Western blot analysis.

### Salt Extraction of Nuclei

Salt extraction of chromatin was performed as previously described ^8^. Cells were harvested from plates prior to protocol and snap frozen on liquid nitrogen and stored at -80 prior to extraction. Frozen cells were thawed on ice and gently resuspended in 1mL hypotonic buffer (10mM Hepes pH 7.9, 1.5mM MgCl2, 10mM KCl, 1mM PMSF, 0.5mM DTT) and incubated 30 minutes on ice, dounced 40 times with a tight pestle and an aliquot was set aside as whole cell extract. Cells were spun at 1500g x 5 minutes and the supernatant removed. Next 400uL of Buffer III.A (10mM Tris pH 7.4, 2mM MgCl2, 1mM PMSF, 5mM CaCl2) was added gently with a wide orifice p1000 tip and 1uL MNase (NEB, 2000 U) was added. Nuclei were incubated for 30 minutes at 37 degrees in a water bath, mixing every 10 minutes. To stop MNase digestion 25uL of ice-col 0.1M EGTA was added to nuclei and an aliquot was set aside as the nuclei fraction. Nuclei were then centrifuged at 400g x 10 minutes at 4 degrees and the supernatant was kept as the MNase fraction. Nuclei were then washed with 400uL Buffer III.B (Same as Buffer III.A, but without CaCl_2_). Chromatin fractions were isolated by adding 400uL of IV.80, incubating for 30 minutes and spinning as above. The supernatant following each spin was saved, and the next salt buffer was added (IV.80, IV. 150, IV.300, IV.600, 10mM Tris pH 7.4, 2mM MgCl_2_, 2mM EGTA, 0.1% Triton X100, with NaCl added per buffer name). All samples were prepared for western blot by boiling in 1X laemmli buffer containing 0.1M DTT.

### Immunofluorescence

RWPE1;LACZ and RWPE1;SCHLAP1 cells were grown on 0.1% gelatin-coated glass coverslips, fixed in 2% paraformaldehyde, blocked in antibody dilution buffer (goat serum PBS) and immunostained for SMARCA4 (abcam, 1:500 dilution) and SMARCB1 (abcam, 1:400 dilution) overnight at 4 degrees in antibody dilution buffer. Coverslips were washed 3 × 5 minutes in PBS and then stained in secondary antibody (goat anti rabbit, Alexa Fluor 568, 1:500 in antibody dilution buffer) for 45 minutes at RT. Coverslips were again washed 3 × 5 minutes in PBS and mounted with Prolong Gold containing DAPI and imaged on a Zeiss Axio Imager 2.

### Growth Assays

2D growth assays were performed by plating 1000 cells/well in 96 well plate and were assayed at 1, 3, and 5 days post plating by Cell Titer Glo (Promega). 3D Growth assays were performed by plating 10 or 100 cells per well in 96-well plates and counting cell growth by Cell Titer Glo 3D (Promega) at 4 days following plating.

### Immunoprecipitation

Nuclear lysates and co-immunoprecipitation were performed as previously described ^10^. Cells were washed with PBS and then centrifuged at 1300 rpm for 10 min at 4 C. Cells were washed with 20 packed cell volumes with hypotonic cell lysis buffer (10 mM HEPES–KOH pH 7.9, 1.5 mM MgCl2, 10 mM KCl, 0.5 mM DTT plus protease inhibitors) and placed on ice for 10 min to swell. Cells were then centrifuged at 1300 rpm for 10 min at 4 C. Cells were dounced with B pestle in 2 packed cell volumes of hypotonic cell lysis buffer. Nuclei were pelleted at 1300 rpm for 10 min at 4 C, washed with 10 packed cell volumes with hypotonic cell lysis buffer and centrifuged at 5000 rpm for 10 min. Extractions were performed twice with 0.6 volume nuclear lysis buffer (20 mM HEPES–KOH pH 7.9, 25% glycerol, 420 mM KCl, 1.5 mM MgCl2, 0.2 mM EDTA, 0.5 mM and protease inhibitors). Lysates were clarified at 14,000 rpm for 10 min at 4 C between extractions. Lysates were diluted with storage buffer (20 mM HEPES–KOH pH 7.9, 20% glycerol, 0.2 mM EDTA, 0.2 mM DTT) to bring final KCl concentration to 150 mM and stored at - 80.

Prior to beginning the IP, we washed protein A magnetic beads three times with PBS + 0.5% BSA at 4 C. We resuspended beads in 400 uL of 1X + 0.5% BSA, then added 4-10ug of antibody and incubated overnight at 4 C. The following day, we thawed the nuclear lysates on ice. Lysates were added to antibody-conjugated beads and incubated overnight. Beads were washed four times with BC-150 (20 mM HEPES–KOH pH 7.9, 0.15 M KCL, 10% glycerol, 0.2 mM EDTA ph 8.0, 0.1% Tween-20, 0.5 mM DTT and protease inhibitors), 2 × BC-100 (20 mM HEPES–KOH pH 7.9, 0.1 M KCL, 10% glycerol, 0.2 mM EDTA ph 8.0, 0.1% Tween-20, 0.5 mM DTT and protease inhibitors), 1 × BC-60 (20 mM HEPES–KOH, pH 7.9, 60 mM KCl, 10% glycerol, 0.5 mM DTT and protease inhibitors). Proteins were eluted from beads using 2X Laemmli buffer with 100 mM DTT for 10 min at 95 C.

### ChIP-seq

ChIP experiments were performed as previously described ^10^. Cells were fixed for 30 min at 4 C in 0.3% methanol-free formaldehyde, quenched for 5 min with 125 mM glycine, washed three times and snap-frozen in liquid nitrogen and stored at - 80 C. Frozen pellets were thawed for 30 min on ice, resuspended each pellet in 1 mL swelling buffer (25 mM HEPES + 1.5 mM MgCl2 + 10 mM KCl + 0.1% IGEPAL CA-630 containing 1 mM PMSF and 1X protease inhibitor cocktail, Roche) and incubated 10 min at 4 C. Cells were dounced 20 strokes with a ‘B’ pestle, and then, nuclei were pelleted at 2000 rpm for 7 min at 4 C. The nuclei were washed with 10 mL MNase digestion buffer (15 mM HEPES pH 7.9, 60 mM KCl, 15 mM NaCl, 0.32 M sucrose) and pelleted at 2000 rpm for 7 min at 4 C. The pellet was then resuspended in 1 mL MNase digestion buffer per 4e7 cells + 3.3 uL 1 M CaCl_2_ per mL + PMSF (1 mM) and protease inhibitor cocktail (1X, Roche) and then incubated for 5 min at 37 C to warm. We added MNase (NEB M0247S 2000 U/uL) at 0.5 uL/1e7 cells and incubated for 15 min at 37 C with agitation. Following digestion, the MNase was chelated using 1/50 volume 0.5 M EGTA on ice for 5 min. We added 1 volume of 2X IP buffer (20 mM TrisCl pH 8, 200 mM NaCl, 1 mM EDTA, 0.5 mM EGTA), then passed the sample for a 21G needle five times and added Triton X-100 to 1% final concentration. The sample was cleared at 13,000 RPM for 15 min at 4°, and chromatin was used incubated with antibody overnight at 4° [SMARCA4 (Abcam ab110641)/SMARCA2 (Cell signaling 11966) / SMARCB1 (Abcam)]. Antibody/chromatin complexes were captured with protein A magnetic beads (Bio-Rad) for 2 h at 4° and washed 5 times with Agilent RIPA (50 mM HEPES pH 7.9/500 mM LiCl/1 mM EDTA/1% IGEPAL-ca-630/0.7% Na-deoxycholate) and once with 10 mM Tris/1 mM EDTA/50 mM NaCl. DNA was eluted at 65 C with agitation using 100 uL 1% SDS + 100 mM NaHCO3 made freshly. Cross-links were reversed overnight by adding 5 uL of 5 M NaCl and incubating at 65 C. DNA was treated with 3 uL RNaseA for 30 min at 37 C and then proteinase K for 1 h at 56 C and purified with Zymo Clean and Concentrator ChIP Kit and quantified using qubit before library preparation (Kapa Hyperprep).

### ChIP-seq Analysis

Reads were aligned to hg38 using bowtie2 ^28^ using the sensitive parameters, and duplicates were removed using SAMtools ^29^. For visualization, bigwig tracks were generated using DeepTools ^30^(version 2.5.2), bamCoverage tool with a binsize of 30 bp and extending fragments to the approximate nucleosome size of 150 bp. Tracks can be visualized using IGV ^31^, and bigwig files are available in GEO Accession number GSE114392.

#### Peak calling

Peaks were called using Macs2 (version 2.1.0 ^32^) using the narrowpeak mode using the following parameters. Qval = 0.05 –keep-dup-all --fix-bimodal –nomodel –extsize 150. Additionally, we filtered the peaks against the ENCODE blacklist regions. Peaks were then annotated for the nearest transcription start site using ChIPPeakAnno ^33^ relative to GENCODE v26.

### ATAC-seq

ATAC-seq was performed as previously described ^14^ with modifications. Briefly, 50,000 cells were harvested and lysed in a buffer containing 0.05% Ipegal CA-630 before transposition with a Nextera library prep kit containing TN5 transposomes. Libraries were amplified using 6-8 PCR cycles to enrich for TN5 products and add indexes and sequenced as paired-end 50bp libraries on a Hiseq 2500. ATAC-seq data are available under GEO Accession number GSE114391.

### ATAC-seq analysis

Nextera adapter sequences were trimmed using trim_galore and reads were aligned to hg38 using bowtie2 with the -X 2000 setting ^28^. We removed any reads mapping to the mitochondrial genome and filtered any reads with a mapping quality less than 20 using SAMtools ^29^. Peaks were called using Macs2 (version 2.1.0^32^) using the narrowpeak mode using default settings and --keep-dup-all. Differential openness was identified use csaw ^34^ with window size 150 and background window size 5000 and an adjusted FDR of 0.05 for the combined windows. Motif analysis was performed using HOMER and comparing the open or closed ATAC sites to the background set of static sites ^35^.

## Data Availability

All data are available from GEO under accession number GSE114394.

## Acknowledgements

We thank members of the Magnuson Lab for helpful comments and discussion. We thank Bernard Weissman for the G401;SNF5 inducible cell line and Arul Chinnaiyan for the SCHLAP1 expression construct. This work was supported by grants to JRR from The University of North Carolina at Chapel Hill University Cancer Research Fund (Tier 1 Development Award) and North Carolina Translational and Clinical Sciences Institute (2KR851610). Additionally, this work was supported by Grants from The National Institute of Child Health and Development (TM 5R01HD036655) and The National Institute of General Medical Sciences (JRR 5F32GM108367).

## Author Contributions

JRR and TM contributed to conceptualization; JRR, KS, JMC took part in methodology; JRR, KS, CJM, and CCS performed investigation; JRR involved in data curation; JR and TM took part in writing—original draft; JRR, KS and TM contributed to writing—review and editing; JRR and TM involved in funding acquisition; JRR and TM carried out supervision. All authors read and approved the final manuscript.

**Supp. Fig. 1.**
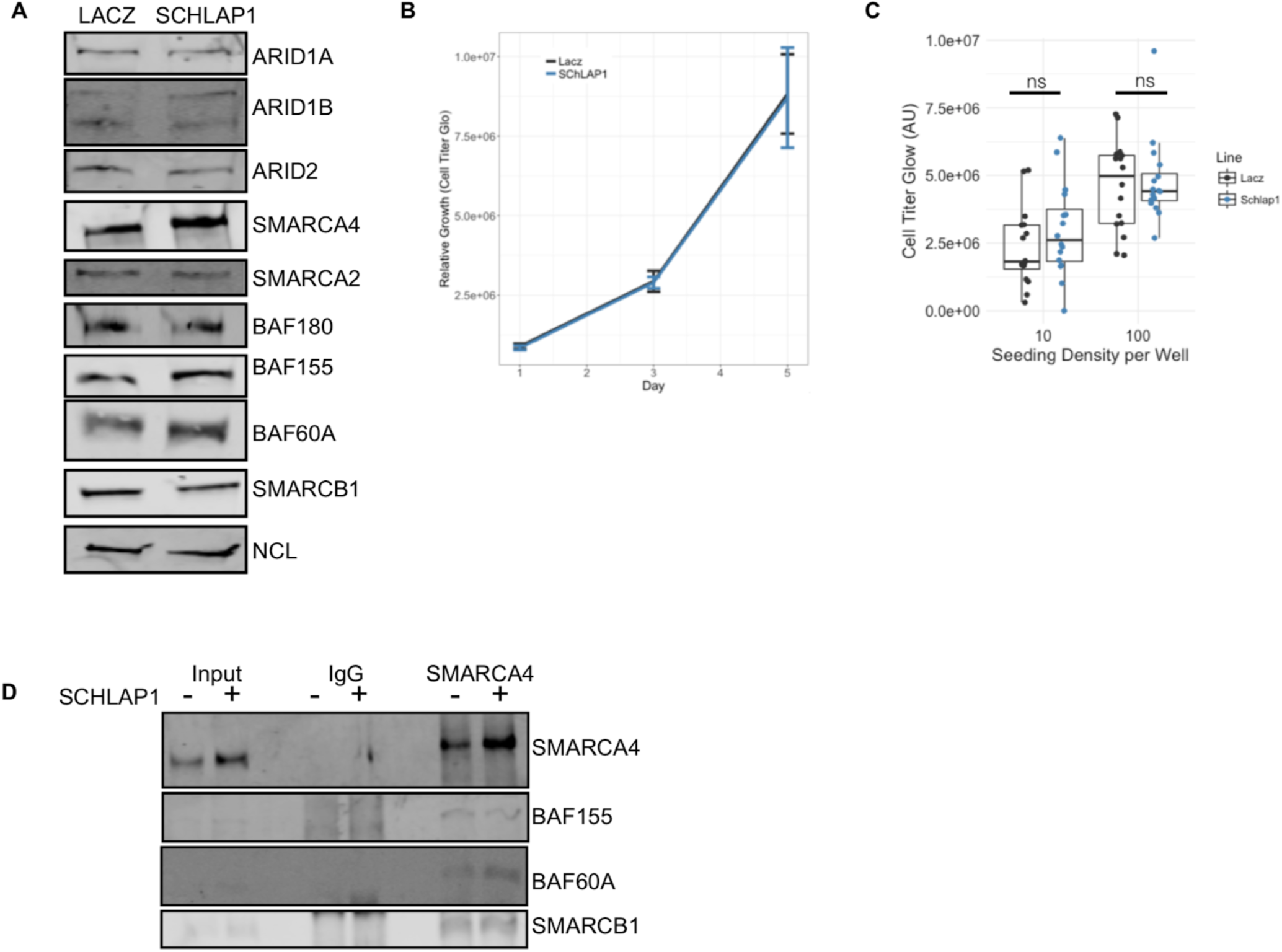
*SCHLAP1* has minimal effects on SWI/SNF composition and cell growth. A. Western blot of SWI/SNF subunits in RWPE1;*LACZ* and RWPE1 ;*SCHLAP1* cells. B. Growth of RWPE1;*LACZ* and RWPE1; *SCHLAP1* measured in two-dimension growth assay. C. Growth assay performed on cell lines in B starting from 10 or 100 cells per well in 96-well low adherent plates. D. Co-immunoprecipitation for SMARCA4 and western blot for other SWI/SNF subunits.

**Supplemental Figure 2.**
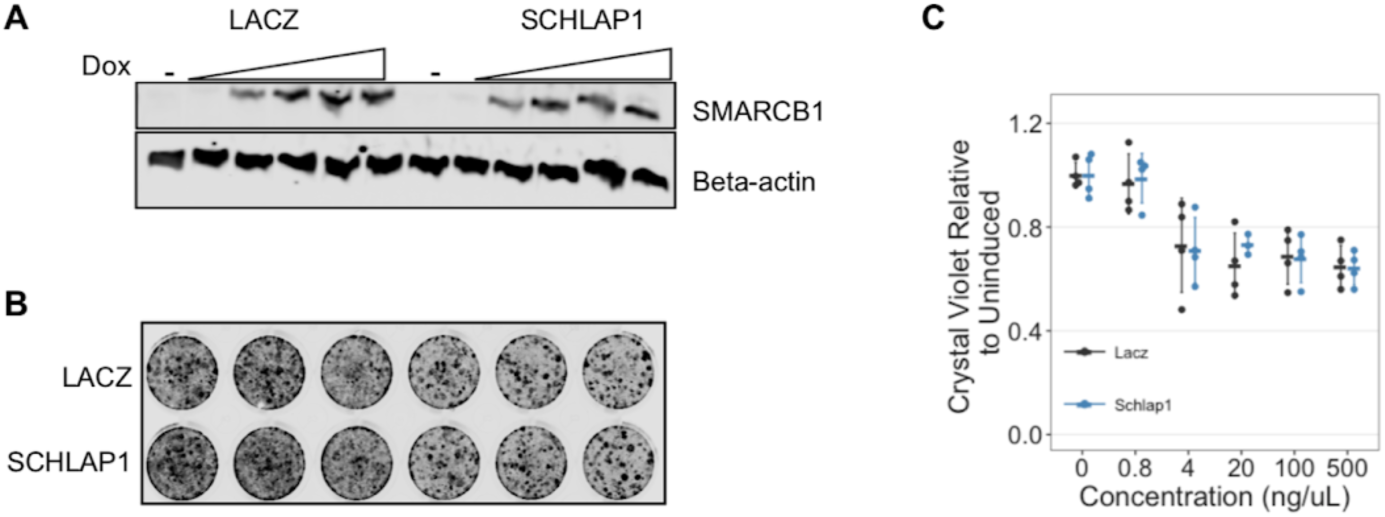
SCHLAP1 does not disrupt SMARCB1 from arresting G401 cells. A. Western blot showing doxycycline induction of SMARCB1 in G401;*LACZ* or G401; SCHLAP1. B. Representative images of crystal violet stained G401 cells. C. Quantitation of images in B (n = 4), using licor imaging.

**Supp. Fig. 3.**
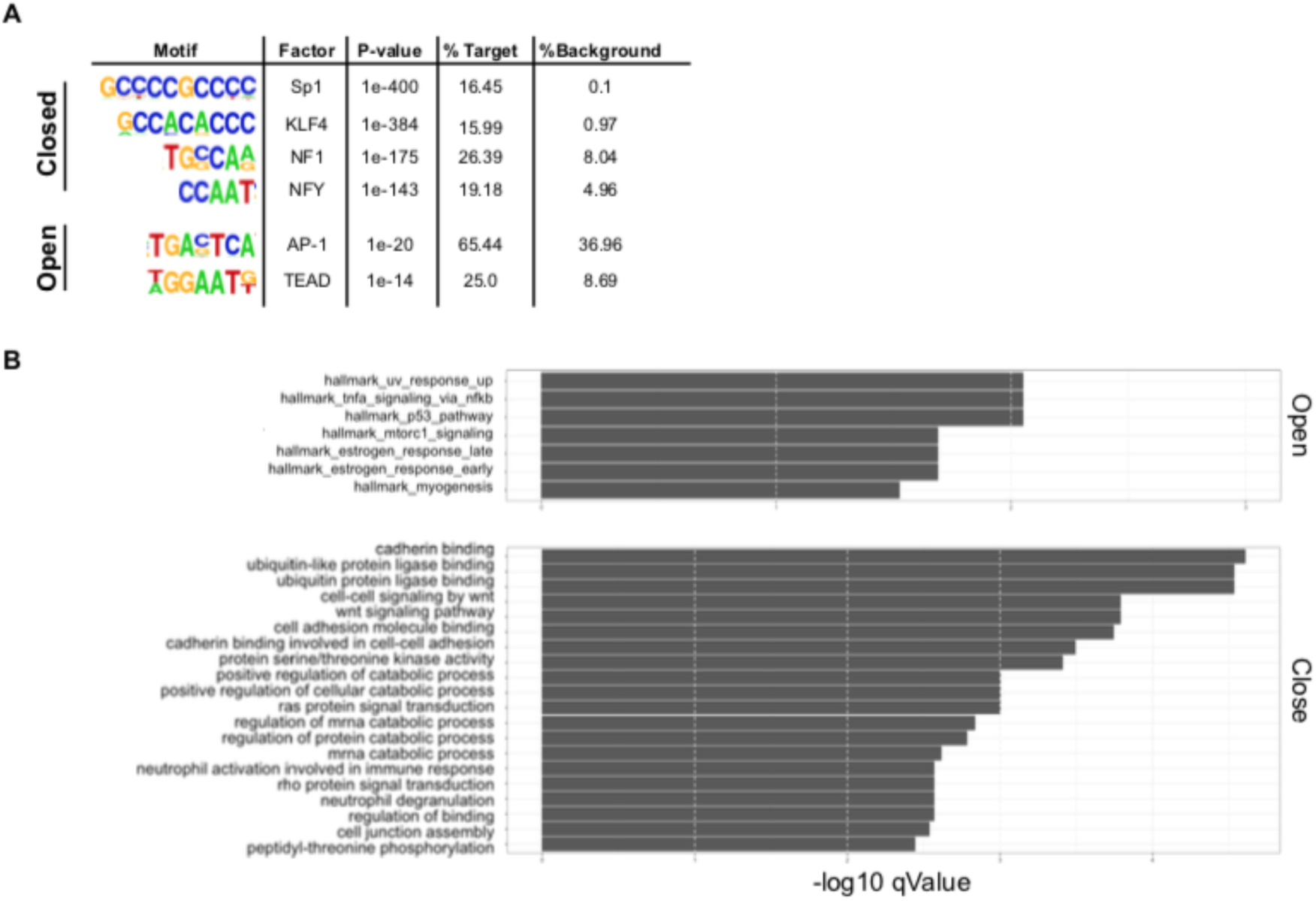
SCHLAP1 induced chromatin changes are associated with specific processes and motifs. A. Enrichment analysis of sites that either open or close following SCHLAP1 expression in RWPE1 cells. B. Motif analysis showing enriched motifs in open compared to closed, or closed compared to open.

**Supp Fig. 4.**
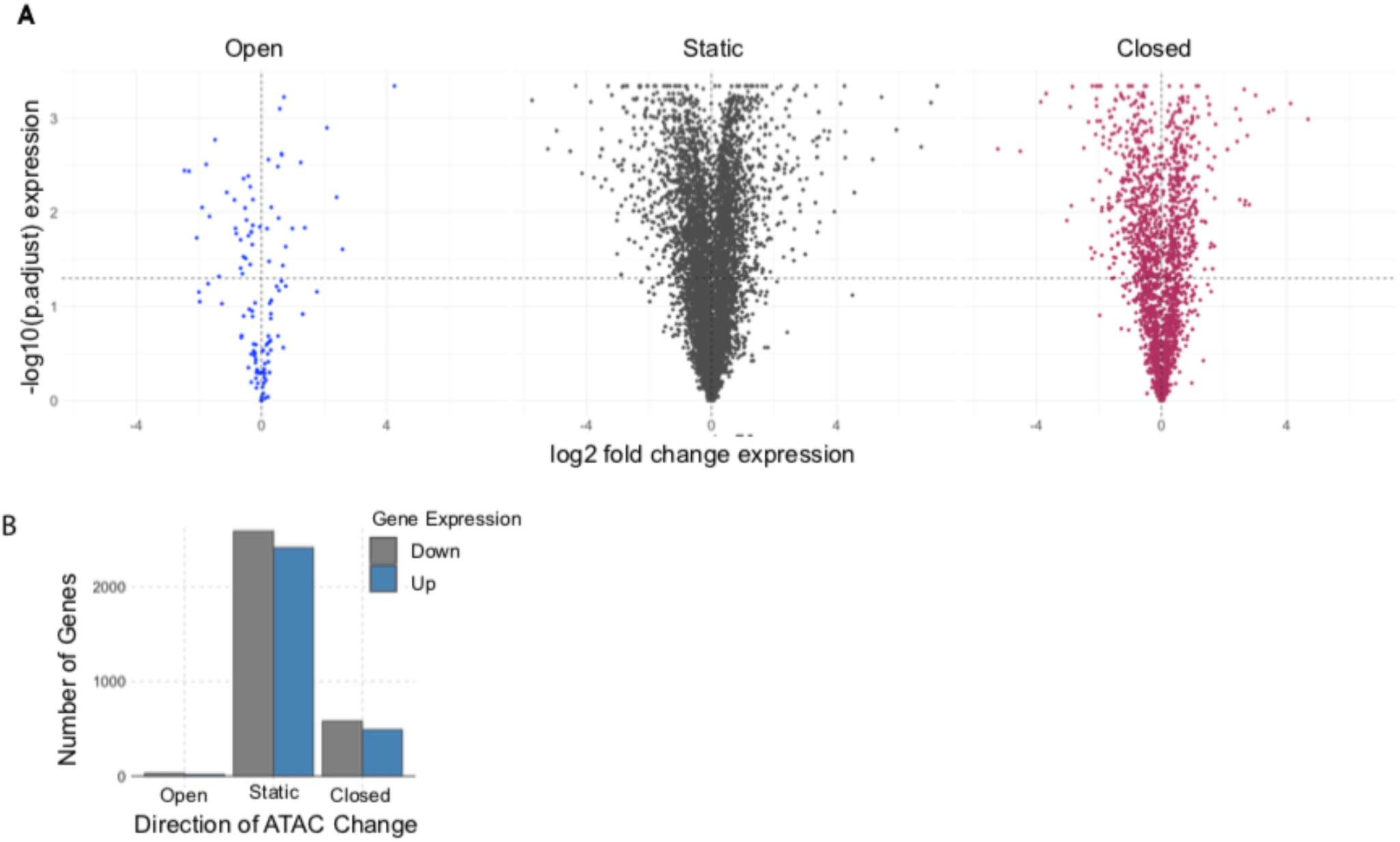
SCHLAP1 induced chromatin changes do not cause concordant gene expression changes. A. Significance and changes in expression of gene expression microarrays performed by Prensner et al. ^1^ on RWPE1;LACZ or RWPE1;SCHLAP1 compared to the altered chromatin regions. Each panel represents the set of regions that either Open, Close, or remain Static in our ATAC-Seq assay. B. Quantification of the number of genes that are significantly differentially expressed and that are associated with an ATAC-Seq peak. These represent those genes that fall below the p-value threshold of 0.05 in panel A.

